# Phase-separating MoSpa2 Complex Organizes Actin Nucleation Center for *M.oryzae* Plant Infection

**DOI:** 10.1101/2024.05.31.596866

**Authors:** Danxia He, Yuanbao Li, Qianqian Ma, Libo Han, Dingzhong Tang, Yansong Miao

**Affiliations:** School of Biological Sciences, Nanyang Technological University, 637551, Singapore; State Key Laboratory of Ecological Control of Fujian-Taiwan Crop Pests, Key Laboratory of Ministry of Education for Genetics, Breeding and Multiple Utilization of Crops, Plant Immunity Center, Fujian Agriculture and Forestry University, Fuzhou 350002, China; Institute for Digital Molecular Analytics and Science, Nanyang Technological University, 636921, Singapore

## Abstract

Polarized actin cable from Spitzenkörper at the hyphal tip fuels filamentous growth in diverse biphasic fungal pathogens. This multi-component complex, featuring the actin nucleator Bni1 and other factors, initiates actin polymerization, guiding biphasic fungal growth and host infection. How dynamic assembly of Spitzenkörper and actin cable are achieved to support filamentous fungi that undergo multistage morphogenesis for host invasion remains unclear, including *Magnaporthe oryzae*, which undergoes multistage morphological transition during rice infection. Here, we identified that the scaffolder MoSpa2 remodeling actin cable networks, in space and time, by assembling the polarisome complex via phase separation, supporting *Magnaporthe oryzae’*s polarized growth. Via N-terminal intrinsically disordered regions (IDRs), MoSpa2 first stimulates actin cable assembly through multivalent interactions with MoBni1 nucleator, and then also creates polarized actin cable bundles by F-actin association and a concurrent inhibition of cofilin-mediated F-actin depolymerization. MoSpa2 mutants exhibit impaired hyphal growth and reduced rice infection, underling its significance. This work elucidates the fundamental mechanisms underlying fungal morphogenesis, offering the potential for targeted interventions in pathogenesis.

## Introduction

The establishment of fungal polarity is a sophisticated process that necessitates the meticulous organization and control of the cellular machinery responsible for polar growth and interspecies communications (Xie & Miao, 2021; Xie *et al*, 2019; Xie *et al*, 2022). The multi-component polarisome complex is central to polarity establishment and directed cell growth in yeast and dimorphic fungal pathogens (Foltman *et al*, 2018; Meyer *et al*, 2008; Moseley & Goode, 2005a). Various components of the polarisome are recruited to the polarized fungal tip, including the scaffolder Spa2 (Araujo-Palomares *et al*, 2009), the actin nucleator Bni1 (Martin Pring, 2003), actin nucleation promoting factors (NPFs), such as Bud6, Aip5, and Pea2 (Xie *et al*., 2019). These elements orchestrate actin cables to facilitate polarized cargo transport and secretion (Steinberg, 2014). During the growth of fungal hyphae, polarisome components combine with secretory vesicles to form a protein-rich Spitzenkörper (SPK) scaffold, promoting filamentous growth (Steven D. Harris, 2005; Zheng *et al*, 2020).

The morphogenetic progression in fungi involves the dynamic remodeling of actin cables, which is orchestrated by efficient actin nucleation activities, actin crosslinking, and polarisome assembly, which spatially and temporally controls elongation (Ghose & Lew, 2020; Ma *et al*, 2021; Sudbery, 2008; Sun *et al*, 2021; Xie & Miao, 2021). The underlying forces that enable host penetration are governed by the assembly status of the polarisome complex. This complex provides essential biochemical activities for polymerizing actin cables both before and after the yeast-to-hyphae transition. To infiltrate the host’s physical barriers, plant fungal pathogens undergo intricate, multistage morphogenesis, which involves the directional transport of virulence factors via the actin cytoskeleton system (Jones & Sudbery, 2010; Prostak *et al*, 2021). An example of a fungus, which does so, is the ascomycete fungus, the *Magnaporthe oryzae* (*M. oryzae*), causing the rice threading blast fungal disease (Wilson & Talbot, 2009). The infection cycle of *M. oryzae* on rice involves various stages, including spore adhesion, germ tube formation, appressorium formation, penetration peg formation, and invasive hyphae growth within the plant cell (Li *et al*, 2017; Liu *et al*, 2017; Shabbir *et al*, 2022). These sequential morphogenic stages produce various protruding structures, facilitating stable colonization, efficient penetration, and rapid filamentous growth for invasion (Doehlemann *et al*, 2017; Meyer *et al*., 2008; Ryder, 2019; Valent, 1996). However, despite these complex morphogenic transitions, our understanding of how polarisome components assemble, organize, and function spatially and temporally to generate polarized actin cables and guide pathogenesis remains limited.

Polarisome complex components are significantly enriched in intrinsically disordered regions (IDRs) (Xie & Miao, 2021; Xie *et al*., 2019; Xie *et al*., 2022), which exhibit high conformational flexibility due to the absence of well-defined folding. This feature facilitates interactions with multiple partners across diverse biological processes (Bah *et al*, 2015; Miao *et al*, 2018). Characterized by dynamic assembly and disassembly, the polarisome is essential for establishing polarity and promoting filamentous growth. During these transitions, IDR-containing polarisome components support dynamic inter- and intramolecular interactions as required, such as during growth transition or stress adaptation (Staples *et al*, 2023; Xie *et al*., 2019; Xu *et al*, 2021). Spa2, the most abundant and scaffolding member of the polarisome complex, remains largely unexplored with regard to its role in regulating dynamic complex assembly, polymerizing polarized actin cables, and facilitating host infection by filamentous fungi.

Our research on *M. oryzae*’s Spa2 (MoSpa2) uncovers its crucial roles in organizing actin nucleation centers for rice infection, by directing polarity establishment and hyphal growth during infection. We have elucidated the molecular mechanisms whereby MoSpa2 directly interacts and co-phase separates with formin to enhance actin nucleation, and also interacts with F-actin for crosslinking. Firstly, MoSpa2 boosts MoBni1-mediated actin nucleation by undergoing co-phase separation, forming a nucleation center at the growing tip where nucleators are enriched. Secondly, MoSpa2 directly binds to both G-actin and F-actin via its N-terminal IDR, thereby initiating F-actin crosslinking through molecular condensation. This process stabilizes the polarized F-actin, directing it towards the hyphal tip for filamentous growth and invasion. Thirdly, F-actin binding by MoSpa2-IDR counteracts MoCof1-mediated depolymerization, further enhancing bundle formation originating from the MoSpa2-MoBni1 nucleation center. Collectively, *M. oryzae* creates a focused actin polymerization center during the invasive hyphal growth stage, a time when organized actin cables are essential for rapid elongation in rice. Our study outlines a spatiotemporally regulated multicomponent polarisome complex condensation during a specific rapid hyphal growth stage of pathogenic fungal infection, thereby shedding light on the spatiotemporally regulated actin remodeling process.

## Results

### Polarisome scaffolder MoSpa2 regulates *Magnaporthe oryzae* life cycle and its rice infection

Given the significant role of *S. cerevisiae* SPA2 as a crucial component in polarisome-mediated polarity establishment(Peter, 2002; Xie *et al*., 2019; Yi-jun sheu, 1998), we were motivated to investigate the MoSpa2’s function in plant fungus *M. oryzae* for its rice infection. We first generated *MoSPA2* loss-of-function mutants (Δ*spa2*) and complementary (CP) line *SPA2/Δspa2* with C-terminal tagged mCherry (see Methods). Genomic integrated Lifeact-GFP was introduced into Δ*spa2* and wild type (WT) as well as CP to monitor actin cytoskeleton in living cells. 4-week-old rice leaves were inoculated by spores of individual *M. oryzae* strains, followed by pathogenesis monitoring after 5 days post infection (dpi) (Fig 1A). In comparison to WT strains, Δ*spa2* exhibited a substantial reduction in disease progression,, which, notably, was restored in the CP strain-infected plants (Fig 1A and B). A similar infection comparison was also conducted on Arabidopsis by analysing *MoSPA2*-dependent disease symptoms and leave lesions (Appendix Fig S1A and B), revealing compromised host infection in Δ*spa2* compared to WT.

**Figure 1.**
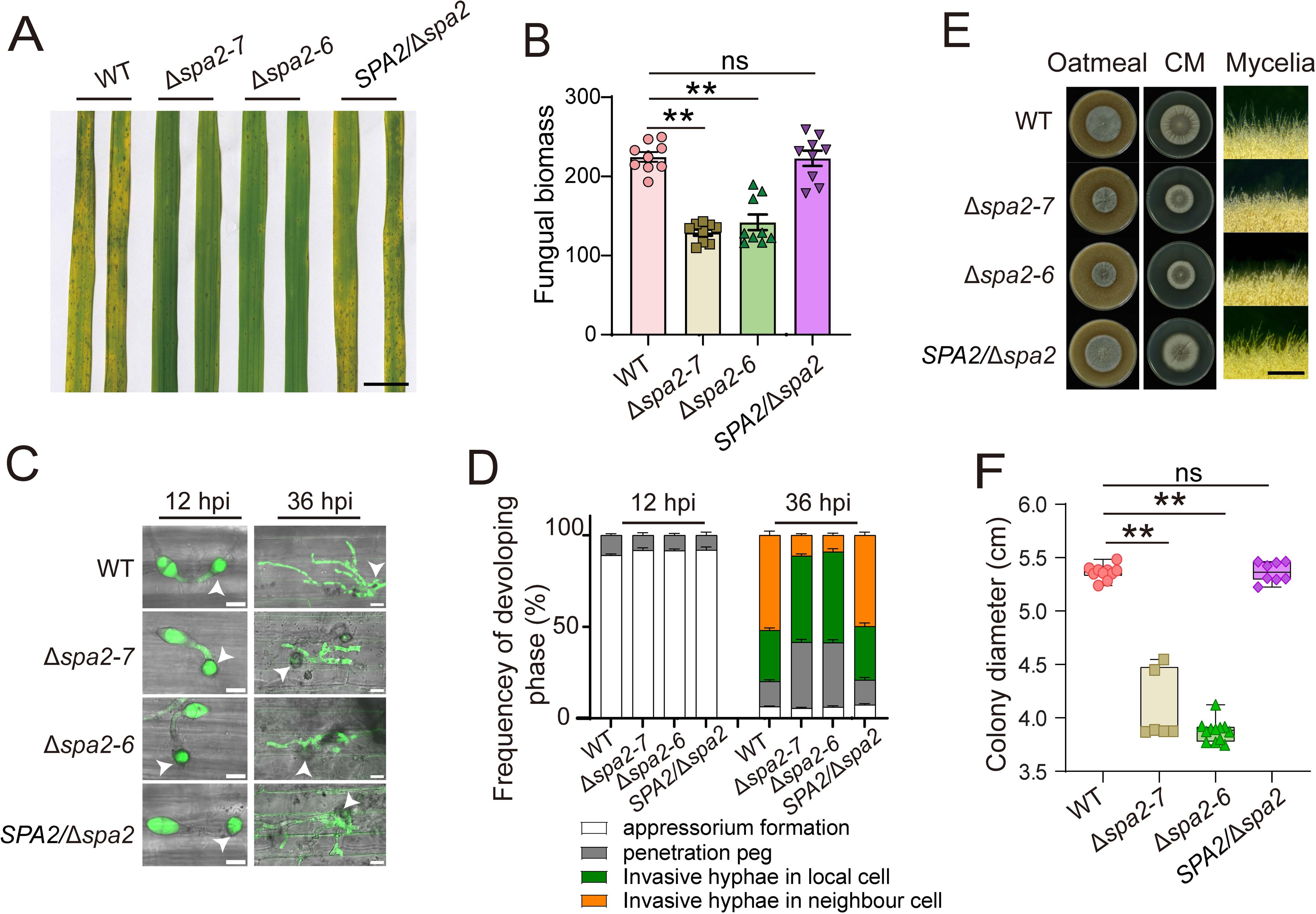
MoSpa2-repressed hyphae growth vitiates disease symptom in rice. **A** Morphology of *M. oryzae* wild type, Δ*spa2* knockout mutants and complementary line pathogenesis in rice. Two independent tests showed similar results, and black bar indicates 2 cm. **B** Quantitative results of *M. oryzae* strains-caused disease. n = 9 leaf disks, (2 x 2 cm^2^ infection leaf area of each). One data point denotes the ratio of gene expression level (MoPot1/UBQ) These DNA was extracted from 2 x 2 cm^2^ infection leaf disk. mean ± SE. Student’s *t*_test was used for statistical analysis relative to WT. ** means *p* < 0.01, ns indicates no significant difference. **C** Morphology at cell level of pathogen infection stage in rice after post 12 and 36 hours post infection (hpi). Two independent tests were performed with similar results. Arrow heads indicate appressoria. White scale bar indicates 10 μm. **D** Quantitative results of different infection stage in (C). n = 396, 366, 392, 394 from 4 leaves in wild type, Δ*spa2-7*, Δ*spa2-6*, complementary line for 12 hpi. n = 517, 590, 542, 536 from 5 leaves in wild type, Δ*spa2-7*, Δ*spa2-6*, complementary line for 36 hpi. mean ± SE. **E** *M. oryzae* growth morphology in oatmeal and cm medium after 7-day growth. Zoom images of mycelia were recorded by confocal. Black scale bar indicates 0.1 cm. **F** Quantification results of *M. oryzae* vegetative hyphae growth length on CM plate from (E). n = 10, 6, 12, 8 values from 5, 3, 6, 4 colonies of wild type, Δ*spa2-7*, Δ*spa2-6*, complementary line. Every each colony was measured 2 times with vertical and horizontal diameter. One-way anova was used for statistical analysis. ** means *p* < 0.01. ns means no significant difference.

We further applied spores of each *M. oryzae* strain onto rice sheath leaves and monitored their progression at specified times (Fig 1C and D; Appendix Fig S1C). Within the initial 12 hpi, more than 85% appressoria was formed by all WT, Δ*spa2,* and CP strains, with no discernible differences (Fig 1C and D). a notable increase in invaded hyphae (IH) was observed in rice sheath cells for WT and CP strains, while *Δspa2* exhibited compromised formation. In the WT and CP strains, IH exhibited burst events into neighboring cells, accounting for 51.9% and 49.8%, respectively. In contrast, only about 10% of IH from *Δspa2* branched into the second host cell (Fig 1C and D). IH of *Δspa2-7* and *Δspa2-6* predominantly remained in the primary infected cells, with rates of 47.1% and 49.6%, respectively, without penetrating another cell for a second time (Fig 1D). This result suggests a sustained penetration into the multicellular walled-plant host, with Spa2 playing a crucial role in facilitating this penetration. A similar reduction in penetration into second neighbor cells was also evident in *Arabidopsis* infection by Δ*spa2* (Appendix Fig S1D and E). Δ*spa2* showed exhibited substantial retention around the appressorium stage during *Arabidopsis* infection.

To pinpoint the specific developmental stages of *M. oryzae* where Spa2 and actin plays a pivotal role in morphogenic transitions for infection—such as conidia, germination, and appressorium formation—we conducted a comprehensive examination. Fungal morphogenesis, along with the actin cable marker Lifeact, was tracked and analyzed without involving the host. Surprisingly, Δ*spa2* strains exhibited no noticeable alterations in the morphogenesis of the mentioned stages or in actin patterns compared to the WT (Appendix Fig S2A-C). However, a distinctive disruption in filamentous growth was observed in Δ*spa2*, unlike the WT and CP strains (Fig 1E). This disruption was evident in both conidia production on oatmeal medium (OMA)(Molinari & Talbot, 2022; Que *et al*, 2020) and vegetative hyphae growth on complete medium (CM) (Jacob *et al*, 2015) (Fig 1E and F). Notably, *Δspa2* exhibited significantly slower hyphal growth compared to the WT (Fig 1F). Despite this, no apparent defects were observed in the growth tip morphology of vegetative hyphae (VH) (Appendix Fig S2D and E), indicating impaired polar growth rather than a disruption in polarity establishment and hyphae tip morphogenesis.

### MoSpa2 organizes an actin polymerization center at the growing hyphal tip

The polarized growth of filamentous fungal hyphae relies heavily on the formation of polarized actin cable bundles. To investigate the role of MoSpa2 in regulating actin cable polymerization and organization in *M. oryzae*, we initially examined a *M. oryzae* line that expresses MoSpa2-mCherry and Lifeact-GFP across various morphological stages. Our analysis revealed a high degree of colocalization between MoSpa2 and Lifeact signals during the conidia and germination tube stages, but not during appressorium formation (Appendix Fig S2F). This suggests both coupled and independent functions of Spa2 and actin filaments in different morphogenetic stages. In the context of hyphae growth, polarized actin cables were observed to polymerize from a highly enriched Spa2 center at the vegetative hyphae (VH) tip (Appendix Fig S2F). This indicates a Spa2 complex-centric formation of actin cables specifically for hyphae growth.

We subsequently investigated the function of MoSpa2 in actin cable polymerization in VH through super-resolution living-cell imaging. In comparison to WT, the actin cable structure in Δ*spa2* was highly disrupted, with either more fragmented or thinner cables (Fig 2A-C). The decreases in actin cable density and skewness (Ma *et al*., 2021; Zhu *et al*, 2017) (Fig 2D and E) were observed, reflecting changes in total actin cable occupancy and individual actin cable filament bundling in VH, respectively. These actin cable defects in Δ*spa2* lines were largely restored in CP. Conversely, due to the compromised cables in Δ*spa2*, there was an increase in the Lifeact-labelled patch pattern (Fig 2C), since a single G-actin pool was shared by actin patches and cables (Miao *et al*, 2013; Yansong Miao, 2016), underscoring the crucial role of MoSpa2 in actin cable formation. It’s worth noting that deletion of Spa2 in budding yeast did not cause a noticeable defect in actin cable formation during polarized growth (Appendix Fig S3A and B), suggesting unique roles for MoSpa2 in the filamentous growth of *M. oryzae.* Strikingly, a clear loss of cable initiation centre at the hyphae tip was observed in Δ*spa2* strains compared to WT and CP lines (Fig 2F; Appendix Fig S3C), implying a function for MoSpa2 in organizing an actin cable polymerization centre.

**Figure 2.**
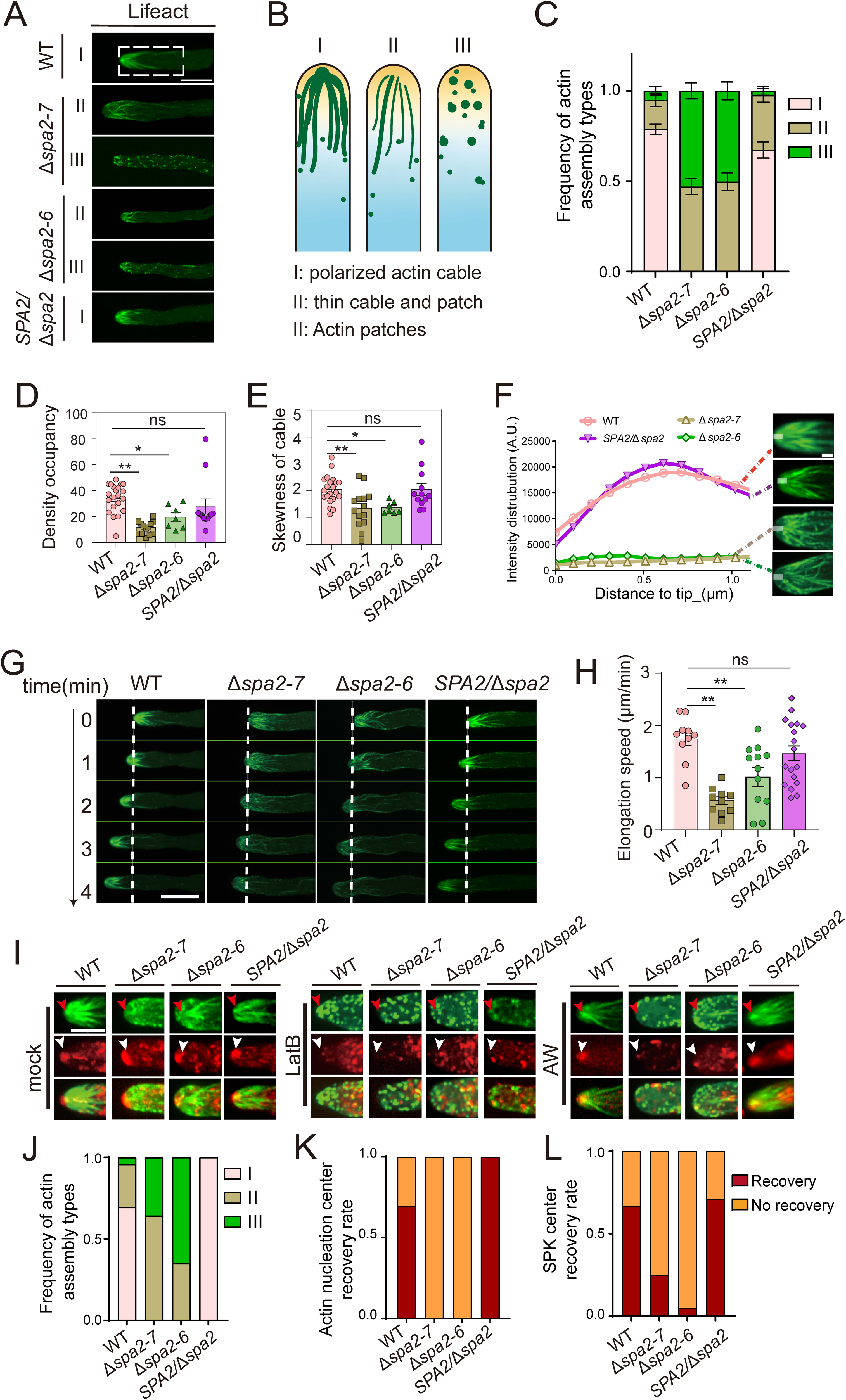
MoSpa2 maintains actin cable stability and organizes actin cable nucleation centre. **A** Living-cell images of *M. oryzae* strains expressing Lifeact-GFP. White box indicates ROI with 100*200 pixels, which is the area for quantification data generation below. White scale bar indicates 10 μm. **B** Classification of actin filament patterns in (A). **C** Quantitative analysis of F-actin in (A). Value indicates means ± SE. n = 26 cells from 4 biological replicated tests in *SPA2/*Δ*spa2* and n = 51, 72, 71 cells from 6 biological replicated tests in wild type, Δ*spa2-7*, Δ*spa2-6.*. **D F**-actin density between multiple strains based on ROI in (A). n = 21, 14, 7, 12 fungal cells in wild type, Δ*spa2-7*, Δ*spa2-6* and *SPA2/*Δ*spa2*. means ± SE. One-way anova was used for statistical analysis. ** means *p* < 0.01. ns means no significant difference. **E F**-actin skewness between multiple strains based on ROI in (A). n = 21, 14, 7, 12 fungal cells in wild type, Δ*spa2-7*, Δ*spa2-6* and *SPA2/*Δ*spa2*. means ± SE. One-way anova was used for statistical analysis. * means *p* < 0.05, ** means *p* < 0.01. ns means no significant difference. **F** Actin cable profile at *M. oryzae* vegetative hyphae tip within 1.326 μm. Representative images of each strains are listed at right side. White scale bar cross tips means region for gray intensity curve production. White scale bar indicates 1.5 μm. **G** Living-cell images with time-lapse of *M. oryzae* vegetative hyphae growth speed. White bar indicates 10 μm. **H** Quantitative hyphae elongation analysis from (G). One-way anova was used for statistical analysis. ** means *p* < 0.01. ns means no significant difference. Value = mean ± SE. n = 10, 10, 12, 18 cells in wild type, Δ*spa2-7*, Δ*spa2-6* and *SPA2/*Δ*spa2* from three independent biological replicates. **I** Representative super resolution images with z-stack (0.2 μm as interval) of vegetative hyphae tip. 100 nM LatB was applied for treatment 20 minutes. AW means after washing of LatB 20 minutes. Green channel means Lifeact-GFP indicating actin cable with red arrowheads indicating nucleation center. Red channel means FM4-64 dye indicating membrane vesicles with white arrowheads indicating SPK centre. White scale bar indicates 5 μm. Two independent tests (Three independent tests for WT) indicates similar results. **J** Quantification of actin assembly types after LatB wash in (I). n = 22 cells from 3 independent tests in wild type, and n = 11, 15, 7 cells from 2 independent tests in Δ*spa2-7*, Δ*spa2-6* and *SPA2/*Δ*spa2*. **K** Regeneration efficiency of actin nucleation centre after LatB wash in (I). n = 22, 11, 15, 7 cells in wild type, Δ*spa2-7*, Δ*spa2-6* and complementary line. **L** Regeneration efficiency of SPK centre after LatB wash in (I). n = 22, 11, 15, 7 cells in wild type, Δ*spa2-7*, Δ*spa2-6* and complementary line.

To evaluate how actin cable defects impact VH growth in Δ*spa2*. We conducted a time-lapse imaging of growing hyphal and measured the elongation speed. In comparison to WT and CP, Δ*spa2* exhibited slower growth in either cells with compromised polar cables (Fig 2G and H; Movie EV1) or largely disrupted cables (Appendix Fig S3D and E; Movie S1). However, these defects in actin cable formation in Δ*spa2* did not appear to result from a disruption of the Spitzenkörper (SPK), a membrane-vesicle-rich centre for polar hyphal growth (Jones & Sudbery, 2010). This is because the FM4-64 dye, which stains membrane vesicles, was still able to indicate an SPK centre at the hyphae tip of Δ*spa2,* if the actin cable is only partially disrupted (Fig 2I). Nonetheless, in Δ*spa2* with high cable disruption or WT treated with a low concentration of latrunculin B (LatB) that only disrupts actin cable but not actin patches, a polarized SPK at the tip was no longer visible (Fig 2I; Appendix Fig S3F). Interestingly, following LatB washout, the reformation of polarized actin cables was observed in WT and CP, along with SPK reformation, whereas Δ*spa2* showed delayed reformation of both polarized actin cables and SPK (Fig 2I-L). These cell biology findings collectively suggest that the generation of the polarized cables from hyphae tip by MoSpa2 is crucial for organizing SPK and enabling polarized hyphal growth.

### MoSpa2 N-terminal IDR guideds molecular condensation

To investigate how the MoSpa2 domain regulates actin polymerization and stabilization, we conducted a series of *in vitro* biochemical experiments using various recombinant MoSpa2 variants. Employing the E. coli expression system, we generated four distinct MoSpa2 variants and the full-length protein, guided by the Alphafold2 prediction and residue signature analysis (Liang *et al*, 2023) (Fig 3A and B; Appendix Fig S4A and B). The developed protein variants are MoSpa2(1-747), MoSpa2(1-338), MoSpa2(339-747), and MoSpa2(748-951) (Appendix Fig S4C). Notably, MoSpa2(1-338) exhibits a high degree of intrinsic disorder and is rich in charge blocks and Pi-Pi interactions, as revealed by NCPR, FCR, and Pi-Pi analysis (Appendix Fig S4B). These characteristics often play a role in facilitating multivalent interactions that contribute to molecular condensation (Han *et al*, 2023; Miao *et al*., 2018; van der Lee *et al*, 2014; Vernon *et al*, 2018).

**Figure 3.**
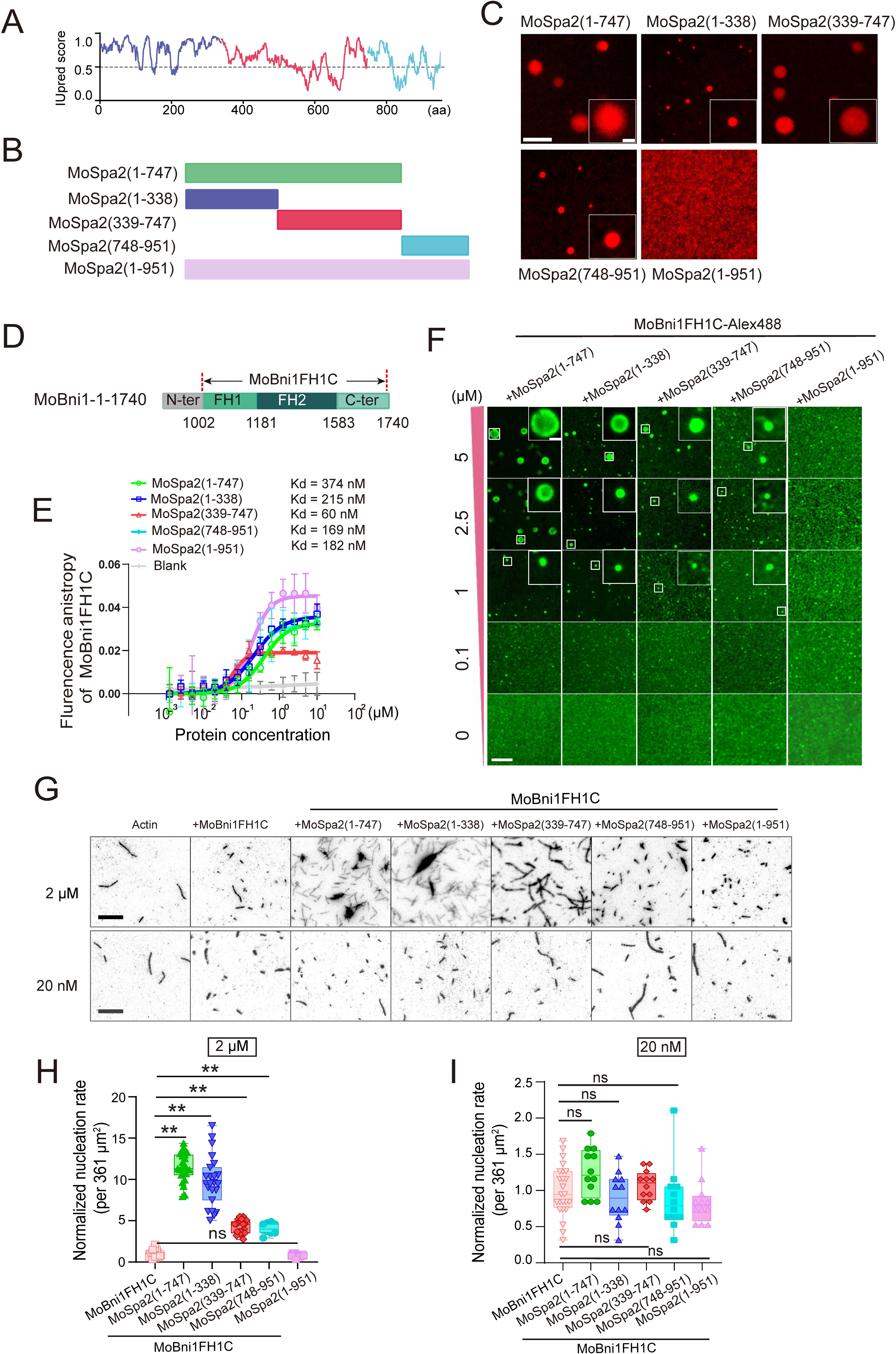
MoSpa2 is phase separating by N-terminal domain *in vitro* and accumulates MoBni1 local concentration to manipulate actin nucleation *in vitro*. **A** IUpred score indicates intrinsic disordered region (IDR) of MoSpa2. Gray dash line at 0.5 indicates the IDR threshold. B Schematic figure for various truncates of MoSpa2 for generating recombinant proteins. **C** Representative images of the phase separation from 5 μM MoSpa2 variants containing 10% Alex-647-labelled protein under 150 mM NaCl. White bar means 5 μm in field region while 1 μm in zoom images. **D** Schematic graph of biochemical active MoBni1 truncating variant, MoBni1FH1C (1002-1740 aa). **E** Anisotropy assay of the interaction between MoBni1FH1C and truncated MoSpa2. Data was fitted with Hill-slope, curve means average with SD of 6 replicates (4 replicates in blank). Blank means 60 nM MoBni1FH1C-Alex488 only. **F** MoSpa2 truncates protein percolation with MoBni1FH1C. 100 nM MoBni1FH1C with 10% MoBni1FH1C -Alex488 was inoculated with different MoSpa2 truncates at different dose under 150 mM NaCl for 5 minutes before imaging. White scale bar means 5 μm. White scale bar in zoom images means 1 μm. **G** Representative TIRF images at 7 minutes after actin polymerization using 2.5 nM MoBni1FH1C and 20 nM or 2 μM MoSpa2 truncates with 0.5 μM G-Actin. Black scale bar indicates 5 μm. **H** Quantitative comparison of actin nucleation efficiency with 2 μM MoSpa2 truncates based on (G). Actin seeds number at 5 minutes were recorded as per ROI at 361 μm^2^, seeds number was normalized to MoBni1FH1C to generate the nucleation efficiency. n = 24, 24, 24, 24, 12, 12 ROIs from two independent test in samples from left to right. One way anova with multiple comparison was used for statistical analysis. ** means *p* < 0.01 **I** Quantitative comparison of actin nucleation efficiency with 20 nM MoSpa2 truncates on (G). Actin seeds number at 5 minutes were recorded as per ROI at 361 μm^2^, seeds number was normalized to MoBni1FH1C to generate the nucleation efficiency. n = 24, 12, 12, 12, 12, 12 ROIs from two independent test in samples from left to right. One way anova with multiple comparison was used for statistical analysis. ns means no significant difference.

Given MoSpa2’s crucial role in structuring a nucleation centre at the growing tip *in vivo*, and its sequence signatures indicating surface charge block- and coiled-coil (CC) oligomerization-mediated phase separation through Pi-Pi interactions, we explored their potential for homotypic demixing across varying protein concentrations and ionic strengths. Notably, MoSpa2(1-747), MoSpa2(1-338), MoSpa2(339-747) and MoSpa2(748-951) all exihibited phase separation dependent on protein concentration and ionic strength but not full-length MoSpa2(1-951) (Fig 3C; Appendix Fig S4D-H). Among the phase variants, MoSpa2(1-338) displayed greater fluidity than the other three variants in the fluorescence recovery after photobleaching (FRAP) assay (Appendix Fig S4I-K; Movie S2). This suggests that MoSpa2(1-338) possesses more dynamic self-interaction compared to the other three variants. Interestingly, full length MoSpa2(1-951) exhibited a diffuse status (Fig 3C), hinting that MoSpa2(748-951) might autoinhibit MoSpa2(1-747) properties for condensation. These results imply that MoSpa2 undergo self-regulation though associations with different variants, eventually coordinating with other polarisome components to regulate *M. oyzae* hyphae polarity growth.

### MoSpa2 promotes MoBni1 mediated actin nucleation via MoSpa2 N-terminus-based multivalent interactions

The involvement of MoSpa2 in actin polymerization *in vivo* and its multivalent capability prompted us to investigate whether MoSpa2 directly regulates actin assembly or relies on the well-known formin nucleator. Initially, using the total internal reflection fluorescence (TIRF) actin assay, we observed that MoSpa2 variants do not induce actin nucleation or promote actin elongation at a concentration of 20 nM (Appendix Fig S5A-C; Movie S3), a concentration that is sufficient for potent nucleator to function. Furthermore, the bulk pyrene F-actin polymerization assay (Appendix Fig S5D-H) substantiates that a concentration lower than 500 nM MoSpa2 variants has no direct effect on actin polymerization, indicating that MoSpa2 is not a potent actin nucleator. Given these findings, we proceeded to reconstitute MoSpa2-involved actin nucleation with MoBni1, the sole formin isoform in *M. oryzae*, as deletion of MoBni1 is lethal (data not shown).

The C-terminal region of MoBni1 (1002-1740aa, MoBni1FH1C), encompassing the biocatalytic domains of polyP-containing FH1, dimeric FH2, and C-terminal IDR, was characterized using Alphafold and distinct physiochemical motif analysis through the machine-learning phase separation engine MolPhase (Graziano *et al*, 2011a; Liang *et al*., 2023; Ma *et al*, 2022a; Miao *et al*., 2013; Moseley & Goode, 2005b; Moseley & Goode, 2006; Sun *et al*, 2018; Xie *et al*, 2020; Xie *et al*., 2019; Xie *et al*., 2022) (Fig 3D; Appendix Fig S6A and B). Then, we generatd biochemically active MoBni1 variant MoBni1FH1C (Appendix Fig S6C) and examined its functional interplay with MoSpa2 variants. Notably, all MoSpa2 truncating variants were found to directly associate with MoBni1FH1C (Fig 3E), suggesting the presence of multivalent interactions between MoBni1 and MoSpa2 for orchestrating actin polymerization. Subsequently, we investigated their macromolecular assembly at physiologically relevant concentrations and stoichiometry. This involved titrating 100 nM MoBni1FH1C with a gradient concentration of MoSpa2 truncates. Interestingly, demixing droplets were observed when 1 μM of MoSpa2(1-338), MoSpa2(1-747), MoSpa2(339-747), and MoSpa2(748-951) were present (Fig 3F), indicating a complex co-phase separation between them.

We then investigated how the phase separation-mediated multivalent interactions of MoBni1 influenced its actin nucleation activities in the presence of different MoSpa2 truncating variants. Using the TIRF actin polymerization assay with 2.5 nM MoBni1-FH1C, we observed that MoBni1FH1C efficiently nucleated actin assembly and generated short F-actin seeds on its own. Intriguingly, all MoSpa2 truncating variants, except the full length, significantly enhanced the overall actin nucleation efficiency above their critical phase-separating saturation concentration (Fig 3G and H; Movie EV2; Appendix Fig S6D and E; Movie S4). This suggests an enhancement in actin nucleation driven by phase separation. A similar synergized actin nucleation effect of MoSpa2 and MoBni1 was observed using the bulk pyrene F-actin polymerization assay (Appendix Fig S7A). The enhanced actin nucleation resulting from the combination of MoBni1 and MoSpa2 can potentially be explained by three biochemical activities. These include: The increased actin nucleation resulting from the combination of MoBni1 and MoSpa2 can potentially be explained by three biochemical activities: 1) the actin nucleation activity of MoSpa2, 2) a nucleation-promoting factor (NPF) activity of MoSpa2 for formin nucleators (such as Aip5 and Bud6 for ScBni1) (Graziano *et al*, 2011b; Graziano *et al*, 2013; Xie & Miao, 2021; Xie *et al*., 2019; Xie *et al*., 2022), and 3) an augmentation in MoBni1 activities due to MoSpa2-mediated multivalent interactions of formins, as observed with multivalent Arabidopsis formin (Ma *et al*., 2021; Ma *et al*., 2022a). To explore these possibilities, we examined MoBni1FH1C’s activities in the presence of a low concentration of 20 nM MoSpa2 variants. This MoSpa2 concentration at an 8:1 stoichiometry to MoBni1FH1C is insufficient to provide direct MoSpa2-based biochemical activities (Appendix Fig S5) while maintaining weak formin-Spa2 interactions given their submicromolar affinity. In results, 20 nM MoSpa2 variants were unable to enhance MoBni1FH1-mediated nucleation (Fig 3G and I; Movie EV2; Appendix Fig S7B and C; Movie S5).

To explore the lack of discernible effects of full-length MoSpa2(1-951) on nucleation, we proposed the hypothesis that MoSpa2(1-951) adopts a closed conformation in vitro, preventing associative interaction with MoBni1 for condensation. Accordingly, we scrutinized MoBni1FH1C assembly through single-particle analysis by titrating with both low and high concentrations, below and above the critical saturation concentration of MoSpa2 variants, respectively. At a 20 nM concentration, MoSpa2 variants were unable to directly cluster MoBni1FH1C particles (Appendix S7D). Conversely, the condensed MoSpa2 truncating variants recruited MoBni1 into a co-phase-separating complex (Fig 3F, Appendix S7E). However, full-length MoSpa2(1-951) exhibited neither self-condensation nor condensation of MoBni1 (Appendix S7E). These collective findings suggest that while a full length MoSpa2 is capable of associating with MoBni1 at submicromolar concentrations (Fig 3E), a self-multivalent condition is crucial to enable its ability in clustering and activating MoBni1 *in vivo*.

### MoSpa2 stabilizes actin cables via crosslinking F-actin and inhibiting MoCof1-mediated depolymerization

In addition, we also noted a significant F-actin crosslinking effect when two phase separating proteins of MoSpa2(1-747) and MoSpa2(1-338) were present (Fig 3G; Movie EV2). This indicates the pronounced biochemical activities of IDR-rich MoSpa2(1-338) in binding and bundling F-actin, critical for providing stable actin cables for polarized growth. First, we investigated the interaction between MoSpa2 and monomeric actin using non-polymerizable G-actin (NP-G-actin) (Xie *et al*., 2020) in an anisotropy assay. Suprisingly, the N-terminal IDR containing MoSpa2(1-338), MoSpa2(1-747), and full length MoSpa2(1-951) could interact with NP-G-actin, while MoSpa2(339-747) did not. Notably, MoSpa2(748-951) also showed association with NP-G-Actin (Fig 4A). Second, the phase separating MoSpa2(1-747) and MoSpa2(1-338) at 2 μM bound and crosslinked preassembled F-actin (Fig 4B; Appendix S8A and B), while MoSpa2 variants lacking the first 338 amino acids and full length failed to associate with or crosslink F-actin (Fig 4B; Appendix S8A and B). Importantly, at a concentration of 20 nM, neither MoSpa2(1-747) nor MoSpa2(1-338) could crosslink F-actin (Appendix Fig S5A-C). These findings highlight that the multivalent and condensation properties of MoSpa2(1-338) enable F-actin bundling activity through binding to G-actin and F-actin. This function likely supports the polarized polymerization of actin cable bundles, fostering hyphae growth.

**Figure 4.**
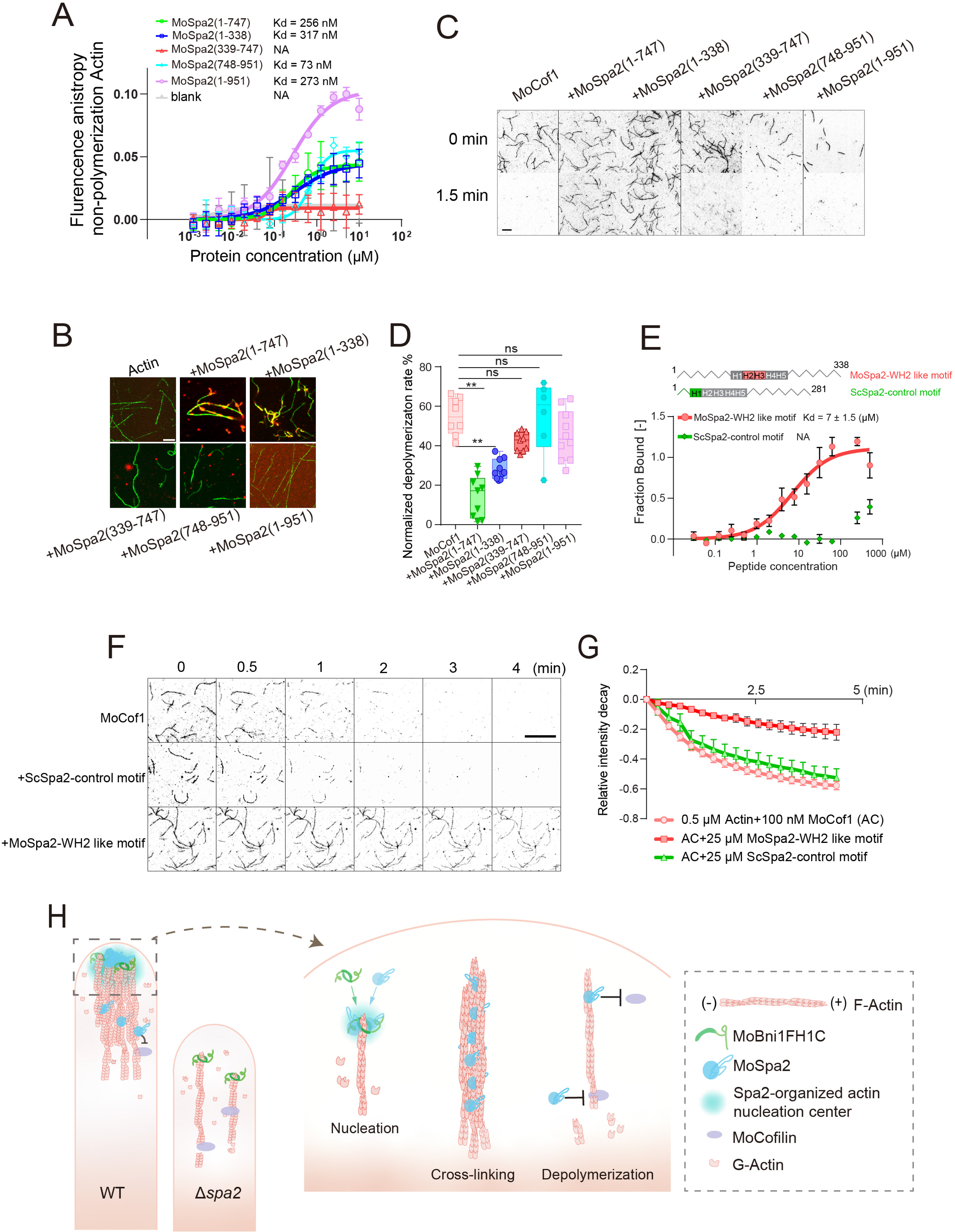
MoSpa2(1-338) contains WH2-like motif that inhibits MoCof1-mediated F-actin depolymerisation. **A** Anisotropy result of non-polymerized G-actin with MoSpa2 truncates. Data was fitted with Hill-slope in prism, curve means average with SD of 6 replicates (4 replicates in blank). Blank means 60 nM MoSpa2 truncates-Alex647 only. **B** Representative images of F-actin-Spa2 sample. Green colour indicates phallodin488-stained actin filaments, red colour indicates truncates of MoSpa2 with 10% Alexa647-labled protein. Actin filaments that polymerized from 2 μM G-actin were diluted 60 times and stained with final concentration of 20 nM Phallodin488 for visualisation. MoSpa2 truncates at final dose of 2 μM. White scale bar indicates 5 μm. **C** Representative TIRF images at indicated time of post 100 nM MoCof1-drived actin depolymerisation in the presence of indicated MoSpa2 truncates at 4 μM. Black scale bar indicates 5 μm. **D** Quantification analysis of MoSpa2 truncates inhibition effects on MoCof-drived actin filaments depolymerization. n = 8, 9, 10, 9, 6, 10 ROIs with 1047.65 μm^2^ in MoCof1, MoSpa2(1-747), MoSpa2(1-338), MoSpa2(339-747), MoSpa2(748-951), 959.36 μm^2^ for MoSpa2(1-951). Relative change in total signal intensity were normalized by comparing to that sample’s first frame where initial depolymerisation start. One way anova with multiple comparison was used for statistical analysis. ** indicates *p* < 0.01. ns means no significant difference. **E** Microscale Thermophoresis (MST) result of binding affinity between MoSpa2-WH2 like peptide and ScSpa2-control motif with non-polymerized G-actin. Value = mean ± SD, n = 6 replicates from two independent test. **F** Time lapse TIRF images of 25 μM MoSpa2-WH2 like peptide effects on inhibition of 100 nM MoCof1-drived actin filaments depolymerisation. Black scale bar indicates 10 μm. **G** Quantification data from (F). Data indicates average with SD from 5 ROIs with per ROI at 1137.7 μm^2^ (528*510 pixels) for Actin and +ScSpa2-control motif, 6 ROIs for +MoSpa2-WH2 like motif in (F). **H** Work model in this study. MoSpa2 and MoBni1 are two main components of polarisome of *M. oryzae*. MoBni1 is an actin nucleation regulator. MoSpa2 is a scaffolder protein responsible for polarisome condensation for actin nucleation centre stability and actin cable formation by crosslinking. Actin cable with stable nucleation centre mediated by MoSpa2 at hyphae tip drives polarity hyphae growth. MoSpa2 as actin assembly regulator at multiple steps including coordination with MoBni1 for actin nucleation, crosslinking actin filaments for actin cable formation and inhabitation on actin depolymerisation factor-drived actin depolymerisation by direct competition of binding with actin.

Next, we endeavoured to unravel the mechanism by which the MoSpa2 N-terminal IDR enabled actin polymerization and stabilization cooperate with effective depolymerization. We delved into the intricate relationship between the N-terminus of MoSpa2 and *M. oryzae* cofilin MoCof1 in actin depolymerization by employing the TIRF actin assay. Recombinant MoCof1 exhibited potent depolymerization activity, reducing preassembled F-actin significantly within 90 seconds (Fig 4C and D; Movie EV3; Appendix Fig S8C and D; Movie S6). Intriguingly, the introduction of MoSpa2(1-747) and MoSpa2(1-338) markedly slowed MoCof1-induced depolymerization, unlike the other MoSpa2 variants and full length (Fig 4C and D; Appendix Fig S8C and D). Furthermore, the MoSpa2(1-338)-mediated deceleration in depolymerization was also evident when ScCof1 from budding yeast was employed (Appendix Fig S8E and F; Movie S7), suggesting a conserved inhibition of cofilin family-mediated actin depolymerization by MoSpa2.

Given the negligible effects of full-length MoSpa2(1-951) on cofilin-mediated depolymerization, we hypothesize suggests that MoSpa2(748-951) may play a role in auto-inhibiting MoSpa2(1-747) containing the 1-388 amino acids against MoCof1. In the presence of MoCof1, we supplemented the reaction with additional MoSpa2(748-951) and MoSpa2(1-747) to compare the F-actin depolymerization rate. While both MoSpa2(1-338) and MoSpa2(1-338)-containing MoSpa2(1-747) inhibited MoCof1-mediated actin depolymerization, the addition of MoSpa2(748-951) to MoSpa2(1-747) seemed to abolish the inhibition of MoCof1 by MoSpa2(1-747) (Appendix Fig S8G and H; Movie S8). This suggests that MoSpa2(748-951) could associate with MoSpa2(1-747), leading to an auto-inhibitory conformation and providing dynamic functionality of MoSpa2 at different *M. oryzae* infection stages where F-actin and MoSpa2 need to couple and decouple (Appendix Fig S2F)."

The role of MoSpa2 in binding, bundling, and competing with cofilin is reminiscent of the phase-separation bacterial effector XopR, which employs a WH2-like domain to counteract cofilin-mediated depolymerization(Dominguez, 2009; Sun *et al*., 2021). Through a meticulous alignment of well-characterized WH2 sequences with helices within MoSpa2(1-338) and drawing insights from the MoSpa2 structure, we identified a potential WH2-like helix, although a conventional LKKV motif is lacking (Appendix Fig S9A). Alphafold prediction of the MoSpa2-WH2-like peptide with actin suggested a 2-helix motif without the conventional LKKV motif for actin binding between subunit 1 and 3 (Appendix Fig S9B). This motif bears resemblance to the 2-helix containing WH2-like motif of XopR (Sun *et al*., 2021). As a negative control, we employed a corresponding helix sequence in the N-terminal IDR of ScSpa2, ScSpa2(1-281), given its inability to associate with F-actin (Appendix Fig S9C). To directly investigate the competition between MoCof1 and the MoSpa2 WH2-like motif, peptides were synthesized, incorporating the ScSpa2-control motif. Microscale thermophoresis (MST) unveiled a distinct binding affinity between MoSpa2-WH2 like peptide and NP-G-actin, with a Kd of ∼6.55 μM (Fig 4E). This peptide inhibited MoCof1-induced actin depolymerization, a trait not shared by the ScSpa2-control motif, mirroring the effects of MoSpa2(1-338) (Fig 4F and G; Movie EV4). These findings collectively illuminate the intricate orchestration that underscores the mechanisms of multivalent and conformation-flexible MoSpa2(1-338) in stabilizing the actin cable efficiently. It adeptly combines actin binding, bundling, and inhibition of MoCof1-mediated depolymerisation.

## Discussion

### Scaffolding protein Spa2 in yeast polarity and rapid fungal hyphal growth

Filamentous fungal morphogenesis and hyphal formation are pivotal for host invasion, promoting physical penetration into host tissues (Dalle *et al*, 2010; Riquelme *et al*, 2018). *Magnaporthe* species, the plant fungal pathogen, undergoes pleomorphic transitions facilitating stepwise plant penetration (Dean *et al*, 2012; Kankanala *et al*, 2007; Wilson & Talbot, 2009). The infection process of *Magnaporthe oryzae* begins with appressorium formation, generating a penetration peg to breach the rice plant’s cuticle and cell wall through the exertion of substantial turgor pressure (Hamer, 1996; Richard j. howard, 1991; Wilson & Talbot, 2009). Then rapid hyphal growth ensues, penetrating neighboring cells and spreading infection (Khang *et al*, 2010). Subsequently, the fungus forms a biotrophic interfacial complex in the invaded cells, where it secretes effectors. These effectors function to suppress the host’s immune responses, eventually leading to the death of the plant tissue (Valent & Khang, 2010). While rapid hyphal growth is critical for a successful plant infection before plant immunity takes place (Doehlemann *et al*., 2017; Jones & Dangl, 2006; Valent, 1996), our understanding of the underlying mechanisms supporting such pathogenic behaviour remains limited.

The formation of polarized actin cables offers a robust pathway for the delivery of secretory vesicles via myosin V, facilitating targeted material conveyance for polarized fungal growth. The condensation of actin nucleators and nucleation-promoting factors is an emerging mechanism guiding directional actin polymerization and cellular morphogenesis (Billault-Chaumartin *et al*, 2022; Ma *et al*., 2022a; Ma *et al*, 2022b; Xie *et al*., 2020; Xie & Miao, 2021; Xie *et al*., 2019; Xie *et al*., 2022). In budding yeast, catalytically active actin regulators, such as the nucleator Bni1 and the formin NPF Aip5, condense in the presence of the multivalent intrinsically disordered region (IDR) and scaffolding protein Spa2. Despite *Saccharomyces cerevisiae* ScSpa2 lacks catalytic activity, it serves as a pivotal scaffolding protein in the polarisome complex, crucial for bud morphogenesis (Xie *et al*., 2019). In fission yeast, the condensation of the formin Fus1 enhances secretion during cell-cell fusion, facilitating concentrated delivery of cell-wall hydrolases (Billault-Chaumartin *et al*., 2022). For filamentous fungi, directional hyphal tip morphogenesis relies on the vesicle-enriched center, Spitzenkörper (Zheng *et al*., 2020). The crowded Spitzenkörper zone generates actin filaments allowing vesicle exchange with the plasma membrane and providing feedback regulation for maintaining Spitzenkörper integrity(Jones & Sudbery, 2010; Zheng *et al*., 2020). Spa2 significantly contributes to the stability of the Spitzenkörper and actin cytoskeletons in filamentous fungi like *Neurospora crassa* (Zheng *et al*., 2020). Deleting the NcSPA-2 gene, encoding Spa2, notably leads to a pronounced reduction in hyphal tip growth rate. These insights underscore Spa2’s multifaceted roles in governing fungal morphogenesis and pathogenicity (Araujo-Palomares *et al*., 2009; Zheng *et al*., 2020).

### MoSpa2 condensation allows focused actin cable formation and Spitzenkörper organization for rapid invasive hyphal growth

The intricate connections among the Spa2 protein, the polarisome, and Spitzenkörper structures remain enigmatic. While the precise biochemical functions and various scaffolding roles of Spa2 within the polarisome and Spitzenkörper are not fully understood, it is acknowledged that Spa2 plays a central role in regulating actin polymerization, crucial for polarized cell growth. Here, in studying the functions of *M. oryzae* MoSpa2 in rice infection, we unveil its multifaceted biochemical and biophysical activities. MoSpa2 directly governs actin nucleation and stabilizes actin cable, promoting rapid hyphal growth, and consequently, effective plant host infection. The initiation of this process involves MoSpa2 establishing an actin nucleation center at the hyphal tip through multivalent interactions with the actin nucleator MoBni1, guiding polarized cable formation towards the rapidly growing tip. Concurrently, MoSpa2 could adopt an open confirmation and directly crosslink F-actin into bundled cables through its N-terminal intrinsically disordered region (IDR)-mediated multivalent interactions with actin, thereby stabilizing the actin bundles for directional filamentous growth. This occurs in conjunction with MoSpa2’s direct competition with the depolymerization factor MoCof1 via a WH2-like motif. The biochemical activities of MoSpa2, both independently and in collaboration with the formin MoBni1, are facilitated by its phase-separating properties via N-terminal IDR. These properties allow the condensation of MoBni1 at the growing tip and MoSpa2’s surface wetting on the F-actin filaments. At present, the mechanisms governing the transition between the open and closed conformations of MoSpa2 are not entirely clear. It appears that MoSpa2 is likely activated timely during the shift to the filamentous hyphae growth stage. In contrast, it maintains a closed, inactive conformation during other stages of the life cycle, refraining from establishing multivalent interactions for the remodeling of F-actin. Our detailed mechanistic exploration of Spa2’s function enhances our understanding of the fundamental processes underpinning the formation and operation of the macromolecular complex at the growing tip of filamentous fungi. This complex comprises the polarized actin cable and concentrated components of the Spitzenkörper. By elucidating how Spa2 facilitates invasive hyphal growth and subsequent infection of the host plant, we gain valuable insights into these intricate biological processes.

## Acknowledgments

We also thank Mr. Yeo Yi Joshua for NP-G-actin purification and Ms. Ng Caixin for the purification of MoSpa2 truncating variants.

## Funding

This study was supported by MOE Tier 2 (MOE-T2EP30121-0015; MOE-T2EP30122-0021 to Y.M), National Research Foundation Singapore NRF-NRFI08-2022-0012; OF-IRG MOH-000955 that is administered by the Singapore Ministry of Health’s National Medical Research Council, and MOE Tier 3 (MOE2019-T3-1-012) to Y.M. This work was also supported by the Distinguished Young Scientists Fund of Fujian Agriculture and Forestry University of China (Kxjq21021) to Y.L.

## Author contributions

Y.M., D.H. and D.T. conceptualized and designed the project. D.H. carried out the majority of the experiments and was responsible for analyzing the data. Y.L. developed the fungal strains and L.H. participated in conducting the pathogen infection assay. M.Q. assisted with the in vitro actin biochemical assay. Y.M. and D.H. prepared the initial draft of the manuscript, with all authors providing valuable contributions to the editing process.

## Competing interests

Authors declare no competing interests.

## Data Availability

This study includes no data deposited in external repositories

## Materials and Methods

### Fungi culture and storage

Fungus used in this study are *Maganoporthe oryzae* wild type strain Y34 (Li *et al*, 2020). Fungus were grown at 28°C on CM medium (yeast extract 6 g/L, peptone 6 g/L, sucrose 10 g/L, agar 15 g/L for plate) ∼ 2-7 days to promote mycelium growth. Fungus were grown at 28°C on oatmeal agar plate (oatmeal 50 g/L, agar 15 g/L) ∼7-11 days for conidia production. Sterilized filter paper (1×1 cm^2^) were used for fungi spore collection and storage. ∼ 5 filter papers were putted onto 4-day-grown fungus oatmeal plate. After ∼6 days, papers covered with fungi were collected and dried by vacuum for long-term storage under - 20℃. *Mospa2* mutants with background of Y34 were generated using the standard one-step gene replacement strategy (Wang et al, 2016; Weld et al, 2006). Briefly, about 1.5 kb DNA fragments flanking the upstream and downstream regions of MoSPA2 gene, were amplified and fused to 5’ and 3’ terminals of a hygromycin resistance gene, respectively. MoSpa2*/Δspa2* complementary line (MoSpa2 gDNA with 1.4kb promoter) fused with mCherry tag was constructed into the pKNTG binary vector, and the DNA sequence was confirmed by sequencing. The primers for all these constructs are listed in Table S1. For the transfer of recombinant DNA fragments or plasmids to *M. oryzae*, 4-day grown Y34 hyphal cultured in liquid CM medium were collected and digested with lysing enzymes (Sigma, L1412) to generate the protoplasts. Then the DNA fragments or plasmids were transformed into *M. oryzae* protoplasts mediated by PEG3350 (Sigma-Aldrich, P4338). Positive *M. oryzae* colones were selected approximately 5 days after transformation (Tang et al., 2015).

### Plant materials and growth conditions

*Arabidopsis thaliana* ecotype Columbia (Col-0) were planted in soil with Vermiculite and pine soil at V:V=1:1 for 4-5 weeks before spore inoculation. The seedlings were grown in a chamber under conditions of 22℃, 10-hour light and 14-hour dark, 65% humidity, 110 μE m-2 s-1. The *japonica* cv. Nipponbare germinated and grown in soil in a growth room with a light/dark cycle of 16/8h at 28°C. 4-week-old rice sheath leaves were used for spore inoculation.

### Plasmid and construction

For transgenic fungi, Lifeact sequence (GGTGTCGCAGATTTGATCAAG AAATT CGAAAGCATCTCAAAGGAAGAAGGCGGCAGCGGC) was introduced into vector pSulPH for Lifeact-GFP fusioned transgenic lines generation. Positive *M. oryzae* colonies were selected by using Chlorimuron-ethyl. All these constructs primers are listed in Table S1.

For protein purification, the MoSpa2(1-747) sequence was synthesised in Twist Bioscience (Sandafold, USA) and introduced into pET28(+) vector (insertion between SacI and HindIII). MoSpa2(1-747) and MoSpa2 short variants were truncated from MoSpa2(1-747)-pET28(+) for protein purification. MoSpa2(1-951) and MoSpa2 (748-951) sequences were synthesised in GenScript (Nanjing, China) and introduced into pNIC28-Bsa4 for protein purification. ScSpa2(1-281) with stop codon was amplified from yeast cDNA and also introduced into pNIC28-Bsa4 for protein purification. MoCofilin (MoCof1) and MoBni1FH1C were amplified from cDNA of *M. oryzae*, and then it was introduced into pNIC28-Bsa4 for protein expression. All primers of truncations are listed in Table S1.

### Pathogen infection

To conduct spray inoculation of conidia, a conidial suspension (the wild type Y34, *Mospa2* mutant and the complemented strain, 1 × 10^5^ conidia/ml) was evenly sprayed onto rice leaves (*Oryza sativa* ssp. *japonica* cv. Nipponbare cultured for 4 weeks) using a sprayer. The infected plants were grown in a growth chamber at 28°C with high humidity in the dark for the initial 24 h, followed by a 12-h/12-h light (20,000 lux)/dark cycle until disease lesions manifested.

For microscopic observation of penetration and expansion of infectious hyphae in rice sheath leaves, 100 μL of the conidial from the Lifeact labeled wild type Y34, *Mospa2* mutant and the complemented strain (1 × 10^5^ conidia/ml) was injected into the inner leaf sheath cuticle cells (*Oryza sativa* ssp. *japonica* cv. Nipponbare cultured for 4 weeks). The the leaf sheath were then incubated under humid conditions at 28℃ and the infection processes of *M. oryzae* in the sheath cell were observed using confocal microscopy (LSM880; Zeiss).

For the pathogen infection on Arabidopsis leaves, 4-5-week-old Arabidopsis plants were used for fungal pathogenesis assay. Conidia from different strains (5 μL at 1×10^5^ spore per m) were applied onto the leaves. After 24 hours, the infected leaves were collected for microscopy imageing of fungi infection. Disease symptom was recorded after ∼3 days.

### Living-cell images and images processing

Fresh fungal mycelium were cultured overnight at 28°C in CM medium with 200 rpm. Refreshed mycelium were mounted onto coverslips for images by spinning disc confocal microscopy coupled with structured illumination microscopy (SDC-SIM). The imaging was performed by a Nikon Ti2 inverted microscope equipped with a Yokogawa CSU-W1 confocal spinning head, a Plan-Apo objective (100 x 1.45-NA), a back-illuminated sCMOS camera (Prime95B; Teledyne Photometrics), and a superresolution module (Live-SR; GATACA Systems). All image acquisition and processing were controlled by MetaMorph (Molecular Device) software. The images were acquired continuously at a 0.2 μm interval for a total range of 20 μm in the z-direction, using 800ms exposure for Lifeact and 600 ms for MoSpa2, and all images were conducted into Fiji for processing. Time-lapse images for hyphae vegetative growth, the image acquisition was done in 0.5 min intervals with 4 min duration. Z-projection was done to each time point frame before composing a final time-lapse file.

### Measurements of Actin cytoskeleton pattern *in vivo*

Fungal tip actin filaments and patch quantification were performed according to previous publications with slight modification (Zhu *et al*., 2017). In brief, area with 200 x100 pixels from hyphae tip was used for actin patch and filaments proportion calculation. Oringinal images were first processed with subtract background at value of 50 and then performed with filter of Gaussian blur at value of 1. Filters processed images were subjected with further analysis. Skewness in vivo (assay area: 200 x 100 pixels) and in vitro (TIRF images were choosed indicated ROIs as present at figure legend) indicating actin cable were determined by using Fiji with plugin of Skewness (Zhu *et al*., 2017). Actin filament density occupancy within hyphae cells was performed by dividing actin filaments signal intensity into cell area (200 x 100 pixels). Actin filaments intensity was identified by setting the same threshold value for visulaized actin filaments intensity among all cells.

### FM4-64 staining and LatB treatment

Fungal transformants expressing Lifeact sequence with fusioned GFP or Spa2-mCherry were stained with FM4-64 (final concentration 5 μM in water). An aliquot of 2 mM stock solution in DMSO of FM4-64 (Cat # 13320, Invitrogen, Carlsbad, CA) was made and stored at -20°C. Vegetative hyphae was first cultured overnight in CM liquid medium and further incubated in a fresh CM solution with FM4-64 dye 5 min for uniform staining before images.

A stock with 5 mM LatB (Sigma, L5288) in DMSO aliquot was prepared and stored at -20°C. Fresh vegetative hyphae was transferred into fresh CM medium with 100 nM LatB for 20 minutes before FM4-64 staining and images. After 20 minutes LatB treatment, vegetative hyphae was washed 5 times by using fresh CM liquid medium before FM4-64 and images.

### Total Internal Reflection Fluorescence assay (TIRF)

TIRF images were conducted according to previous publication (Sun *et al*., 2021), 25 × 50 mm^2^ coverslips (Marienfeld Superior) were cleaned with 20% sulfuric acid overnight and rinsed thoroughly with sterile water. The coverslips were then coated at 70°C water bath with 2 μg/ml biotin-PEG-silane (Laysan Bio Inc.) and 2 mg/ml methoxy-PEG-silane in 80% ethanol (pH 2.0) for overnight. The biotin-coated coverslips were rinsed thoroughly with sterile water, dried in a nitrogen stream, and kept at −20°C for long-term storage. The biotin-coated coverslip was first attached to a plastic flow cell chamber (Ibidi, sticky-Slide VI 0.4), followed by 1 min incubation with HEKBSA buffer (20 mM HEPES pH 7.4, 1 mM EDTA, 50 mM KCl, and 1% bovine serum albumin) and then 60 s incubation with 0.1 mg/ml streptavidin in HEKG10 (20 mM HEPES, pH 7.4, 1 mM EDTA, 50 mM KCl, 10% glycerol (V:V)). After pre-treatment, the flow cell chamber was washed three times by TIRF buffer (10 mM imidazole, 50 mM DTT, 15 mM glucose, 50 mM KCl, 1 mM MgCl_2_, 1 mM EGTA, 100 μg/ml glucose oxidase, 40 μg/ml catalase and 0.5% methylcellulose, 0.3 mM ATP, pH 7.4). Recombinant proteins prepared in TIRF buffer were mixed with 0.5 μM ATP-actin monomer (10% Oregon Green 488 labeled, 0.5% biotin-labeled) before flowing into the chamber. Time-lapse images were acquired at room temperature at 15s intervals for 15 min or indicated interval and duration time when measuring nucleation efficiency and crosslinking effect. Skewness of actin filaments was identified based on above described method. For actin elongation rate quantification, the fast elongation end of each individual filament was traced manually for a period of 2 min of each. Conversion factor of 374 subunits per micrometer of actin filament was used to calculate the elongation rate. For depolymerization, actin filaments were first washed with TIRF buffer 3 times to reduce the actin into constrict dose. Indicated concentration of Cofilin or proteins were flowed into chamber containing constrict concentration actin filaments for depolymerisation. Time-lapse images with indicated duration were acquired at room temperature at 15s intervals.

### Protein sequence prediction

To identify the MoSpa2 sequence characteristics and MoBni1sequence features, analysis of MoSpa2 truncates sites, full length sequence of MoSpa2 and MoBni1 sequence were submitted to MolPhase, which is a novel engine including all below individual traits(Liang *et al*., 2023). ANCHOR (http://anchor.enzim.hu/) for identifying IDR domains. PScore website (http://pound.med.utoronto.ca/~JFKlab/Software/psp.htm) for protein residues Pi-Pi interaction prediction. Sequence residues charge and hydrophobicity parameters were predicted by using website tool CIDER (http://157.245.85.131:8000/CIDER/analysis/). Protein structure was predicated by Alphafold2 with default parameters. (https://colab.research.google.com/github/sokrypton/ColabFold/blob/main/Alphafold2.ipynb)

### Protein percolation assay and Phase diagram

Recombinant MoSpa2 variants were mixed with 100 nM MoBni1FH1C (mixed with 10% Alexa 488-labeled MoBni1FH1C) in buffer (20 mM Hepes, 150 mM NaCl) and incubated at room temperature for 5 min before applying 10 μL on the coverslip for images by SDC-SIM.

For MoSpa2 variants phase diagram, a serial dilution of protein concentration and ionic strength was performed. Volume of 10 μL serial dilution protein in different dose NaCl solution were dropped onto coverslip for 5 minutes before images.

### Pyrene fluorescence assay

Pyrene-labeled actin was purchased from Cytoskeleton Inc. A 10 μM G-actin (5% pyrene actin) was first converted to Mg^2+-^ATP-actin for 5 min on ice before use and then mixed rapidly with various proteins in the G buffer (20 mM Tris-HCl, pH 7.4, 2 mM ATP, pH 7.4, 1 mM CaCl2, 5 mM DTT). The spontaneous actin polymerization was initiated by 10 × KME buffer mix (10 mM MgCl2, 10 mM EGTA, and 500 mM KCl), at a total reaction volume of 120 μL. The pyrene-actin fluorescence signal was monitored in a plate reader Cytation 5 (BioTek, USA) at excitation and emission wavelengths of 365 and 407 nm, respectively. The raw data from 0 to 2400s or 2520s were used to generate fluorescence curve. The fluorescence of each sample was normalized to actin only initial data point. The curve shown in the graph were the average data from at least three biological replicates.

### Anisotropy assay

For measurements of MoSpa2 truncates with NP-G-actin (Xie *et al*., 2020) and MoBni1FH1C, fluorescent proteins were diluted to 60 nM (fluorescein Alexa 488-labeled MoBni1FH1C or Alexa647-labeled MoSpa2(1-747), MoSpa2(1-338), MoSpa2(339-747), MoSpa2(748-951), MoSpa2(1-951) in 1x G buffer for actin association and protein buffer (20 mM Hepes pH 7.4, 500 mM NaCl, 10% glycerol) for protein-protein interaction. The stock of NP-G actin or MoSpa2 truncates were titrated 14 times starting from 10 μM (1:1). The total volume was 24 μL containing 60 nM Alexa-labeled protein. Equal volumes of Alexa-labeled protein and associated protein were incubated for 1h in the dark at room temperature. Alexa 488 fluorescence (485/20, 528/20 nm) or Alexa 647 (635/25, 670/25) was measured by the plate reader Cytation 5 (BioTek, USA).

### Microscale thermophoresis (MST)

Non polymerized G-actin (NP-G-actin) were purified from insect cell and utilized in an affinity test with a WH2-like peptide, employing MicroScale Thermophoresis (MST) as per the standard protocol (Sparks & Fratti, 2019). In brief, MoSpa2-WH2 like peptide (NKARDKLQRLTTVQFLELSTDVYDELNRRF) and ScSpa2-control motif (MGTSSEVSLAHHRDIFHYYVSLKTFFEVT) were first synthesized by GenScript (China). Peptide powder was dissolved in reaction buffer (20 mM Hepes, 150 mM NaCl, 5% glycerol) with 3 mM as stock and kept in -20℃ for long-term storage. For the experiments, the peptides were under serial dilution from 1 mM, to a 2 μM NP-G-actin until 1:1 molar ratio. The mixtures were then gently stirred using a pipette before being transferred into a capillary tube (MonolithTM NT.115). The MST was conducted using a Monolith NT.LabelFree instrument, with the settings adjusted to 20% LED power (red) and 20% IR-laser power, all at room temperature. The resulting data were processed and analyzed using the Mo. Affinity Analysis (x86) software and GraphPad Prism 8.

### Protein purification

For MoSpa2 truncates MoSpa2(1-338), MoSpa2(1-747), MoSpa2(339-747), MoSpa2(748-951) and MoSpa2(1-951) proteins were expressed and purified by using *Escherichia coli* (BL21(DE3) Rosetta T1R). Cells were cultured in 1 L TB medium (24 g yeast extract per liter, 20 g tryptone per liter, 4 mL glycerol per liter, phosphate buffer pH 7.4) containing 50 μg/ml of kanamycin at 37°C to OD_600_ of 1 before induction by 0.5 mM IPTG at 18°C overnight. The cells were harvested by centrifugation at 4°C and 4000× *rpm* (rotor JA10) for 20 min. The pellet was resuspended in binding buffer (20 mM Hepes, pH 7.4 and 500 mM NaCl, 20 mM imidazole, 1 mM PMSF, protease inhibitor cocktail Set III, EDTA free from Thermo Fisher) and lysed by an LM20 microfluidizer (15000 psi). The lysate was clarified by centrifugation at 25,000× *g* for 1 h using rotor JA25.5 (Beckman Coulter). The supernatant was purified by an FPLC AKTAxpress system (GE Healthcare) using Ni-NTA affinity chromatography. The elution fractions containing targeted proteins from gradient elution with increasing imidazole concentrations were pooled and further purified by size-exclusion chromatography using a HiLoad 16/600 Superdex75 column (GE Healthcare) in gel filtration buffer (20 mM Hepes, 500 mM NaCl, 10% glycerol). Proteins were flash-frozen in liquid N2 prior to storage at −80°C in small aliquots.

To obtain monomeric Ca^2+^-ATP-actin with or without labeling with Oregon Green™ 488 iodoacetamide (ThermoFisher) or NHS-dPEG®4-biotin (Sigma), 5 g of rabbit skeletal muscle acetone powder (Pel-Freez Biologicals) was dissolved in 60 ml of G-buffer (5 mM Tris, pH 8.0, 0.2 mM ATP, 0.1 mM CaCl_2_, 0.5 mM dithiothreitol) for 30 min at 4°C. The mixture was filtered through cheesecloth three times to collect the actin-rich extracts in the supernatant. The filtration was repeated three times, and the actin-rich extracts were combined and subjected to centrifugation at 18,000× *g* by rotor JA-25.5 (Beckman). Afterward, the clear actin-rich supernatant was supplemented with 50 mM KCl and 2 mM MgCl_2_ to allow actin polymerization with slow stirring at 4°C for 1 h. To remove actin filament binding proteins, KCl powder was added slowly until a final concentration of 0.8 M and was subsequently stirred slowly for an additional 30 min. The solution was subjected to 45,500× *g* centrifugation for 3.5 h with rotor Ti45 (Beckman) at 4°C to collect the polymerized actin filament. The actin filament pellet was rinsed with 1 ml G-buffer, all the actin filament pellets were transferred to the 10 ml homogenizer with 5 ml G-buffer, and the pellet was homogenized by moving up and down. Actin filament was then dialyzed against 1 L of G-buffer without DTT overnight. The next day, we changed to 1 L of new G-buffer without DTT and kept on dialysis. In the meanwhile, we freshly prepared Oregon Green™ 488 iodoacetamide or NHS-dPEG®4-biotin with high-quality DMSO into a final concentration of 10 mM. The concentration of dialyzed actin was determined by OD_290_ using NanoDrop. Before proceeding to actin labeling, we diluted the actin with cold 2× labeling buffer (50 mM imidazole pH 7.5, 200 mM KCl, 0.3 mM ATP, 4 mM MgCl_2_) to an actin concentration of 23 μM. A 12–15-fold molar excess of Oregon Green™ 488 iodoacetamide/NHS-dPEG®4-biotin stock was added dropwise with very gentle vortex. To allow sufficient labeling, we kept the solution in the dark with aluminum foil and rotated at 4°C overnight. The next morning, we pelleted the labeled actin with Type 50.2 rotor (Beckman) at 111,000× *g* for 3 h. We then collected all the pellets and transferred them to the homogenizer with 5 ml G-buffer. Afterward, the actin was homogenized for 20 times. To further depolymerize the actin, it was dialyzed against 1 L of G-buffer 3 times with 14 hour each time. We then applied the dialyzed actin to centrifugation at 167,000× *g* for 2.5 h with rotor SW55 Ti (Beckman) at 4°C. To further purify actin, the top 2/3 of the supernatant was collected and injected into a GE AKTA FPLC and separated through column HiPrepTM 16/60 SephacrylTM S-300 HR column. The collected labeled actin was dialyzed overnight against G-buffer with 50% glycerol. Small aliquots of actin were prepared and flash-frozen in liquid N2 for long-term storage.

### Fluorescence Recovery after Photobleaching (FRAP)

FRAP experiments were performed on an SDC-SIM with FRAP module, using a 100x oil objective lens. Condensates of MoSpa2 truncates at 5 μM (with 10% Alex647-labled protein) in the condensate buffer (20 mM HEPES, pH 7.4, 150 mM NaCl) for 5 min on coverslip before image. Protein self-assembled lipid droplets were monitored by time-lapse imaging with 200 ms exposure and 15s interval for 5 minutes. The 100% bleaching by 647 nm laser was achieved by five cycles of scanning. For FRAP analysis, we measured mean fluorescence intensities from three regions of interests (ROIs), including one in the photobleached region, one outside of droplet for correcting illumination-caused photobleaching, and one in the background region. Fluorescence recovery was analysed using the plugin FRAP Profiler of ImageJ and plotted by GraphPad Prism 8.

### Protein colocalization with Actin-filaments

Actin filaments were first polymerized from G-actin as above described with slight modification. In detail, 2 μM G-Actin was initiated by 1 × KME buffer mix containing extra 200 μM EGTA and 109.7 μM MgCl_2_ in G buffer. After polymerized in KME buffer for 30 minutes, F-actin was following stained with phollodin488 or phallodin568 for 20 minutes at final concentration of 20 nM. Phallodin-stained F-actin was diluted 60 times following inoculation with 2 μM MoSpa2 variates or 4 μM ScSpa2(1-281) for 20 minutes. Afterword, volume of 20 μL MoSpa2-inoculated F-actin solution was applied onto coverslip for images.

### Statistical analysis

Statistical analyses were performed in GraphPad Prism 8, one-way anova with multiple comparison was used for multiple groups statistical analysis. Unpaired two-tailed student’s *t*_test assuming equal variances was used to determine difference between 2 groups, *p < 0.05; ** p < 0.01; All other significant comparasion with p < 0.001 or p < 0.0001 were grouped into ** p < 0.01. ns means not significant.

## Supplementary table and Expanded movies legends

**Table S1. Primer lists used in this study.**

Primers for *MoSPA2* gene deletion and PCR detection, and for MoSpa2 truncates and related protein purification are listed in this file. Primers working for fungi biomass detection are also listed in Table S1.

**Table S2. Bacteria, fungi strains and plasmids used in this study.**

Bacteria strains used in this work include *Escherichia coli* DH5α and Rossetta for construction vector development and protein purification, respectively. Fungi strains of *M. oryzae* including wild type, mutants and complementary lines were used. Yeast strains used in this study including wild type S288C strain and Δ*spa2* knock out mutants expressed with ABP140-GFP indicating actin cytoskeleton.

**Movie EV1. Hyphae growth tracking via time-lapse images.**

Source images with time lapse (4 minutes in total, 30 seconds interval) for Figure 2G. White scale bar indicates 10 μm.

**Movie EV2. MoSpa2 increases MoBni1 activities for actin nucleation.**

Source images with time-lapse for Figure 3G. Scale bar means 5 μm. Indicated MoSpa2 truncates and MoBni1FH1C were added together into 0.5 μM G-Actin for polymerization. TIRF images were recorded per 15 seconds with duration of 10 minutes and 7 minutes for 2 μM MoSpa2 and 20 nM MoSpa2, respectively.

**Movie EV3. MoSpa2 truncates with MoSpa2(1-338) inhibit MoCof1-drived F-actin polymerization.**

Source data for Figure 4C. Scale bar indicates 10 μm. TIRF images with time 4.15 minutes duration were recorded at 15 seconds interval.

**Movie EV4. MoSpa2(1-338) with WH2-like motif prevents MoCof1-drived actin filaments depolymerization.**

Source images with time lapse (4.15 minutes duration with 15 seconds interval) for Figure 4F. Black scale bar indicates 10 μm. MoSpa2-WH2 like motif and ScSpa2 control motif at 25 μM, MoCof1 at 100 nM and G-actin at 0.5 μM.

## Supplementary Figure and Supplementary Movie Legends

**Figure S1. MoSpa2-repressed hyphae growth vitiates disease symptom in Arabidopsis (related to Fig. 1).**

**A.** Morphology of *M. oryzae* wild type, Δ*spa2* knockout mutants and complementary line pathogenesis in Arabidopsis. Three independent tests showed similar results, black scale bar indicates 1 cm.

**B.** Quantitative results of *M. oryzae* strains-caused disease. Disease symptom value means every lesion area relative to that half leaf area. n = 9, 5, 7, 8 measurement values from 6, 5, 6, 6 leaves in wild type, Δ*spa2-7*, Δ*spa2-6*, complementary line, respectively. Values means average and SE. Student *t*_test was used for statistical analysis relative to WT. * means *p* < 0.05. ns means no significant difference.

**C.** Schematic graph of *M. oryzae* infection and growth stages.

**D.** Representative images at cell level of pathogen infection stage in Arabidopsis after 22 and 48 hours infection. White scale bars indicate 10 μm.

**E.** Quantitative results of *M. oryzae* different infection stage in (D). n = 41, 32, 41, 34 cells in wild type, Δ*spa2-7*, Δ*spa2-6*, CP-infection samples at 22 hours; n = 36, 15, 34, 37 cells in wild type, Δ*spa2-7*, Δ*spa2-6*, CP-infection samples at 48 hours. Values means average and SE.

**Figure S2. MoSpa2 shows no obvious effects on morphology of conidia, germination tube, appressorium and vegetative hyphae tip size. (related to Fig. 1).**

**A.** Spores of wild type and Δ*spa2*. *Z*-stack images with 0.2 μm step were processed for images. White bar means 15 μm.

**B.** Germination tube of wild type and Δ*spa2*. Z-stack images with super resolution with 0.2 μm step were processed for images. White bar means 10 μm.

**C.** Appressorium of wild type and Δ*spa2*. Z-stack images with super resolution with 0.2 μm step were processed for images. Black scale bar means 5 μm.

**D.** White field of vegetative hyphae tip among wild type, Δ*spa2*. Black scale bar means 5 μm, vertical black bar means the area for tip dimeter measurement.

**E.** Quantification data generated from (D). n = 10, 13, 9, 18 in wild type, Δ*spa2-7*, Δ*spa2-6* and complementary line. Student *t*-test was used for statistical analysis, ns means no significant difference.

**F.** MoSpa2 localization at different growth stage of *M. oryzae*. MoSpa2 is tagged with mCherry and Lifeact was fusioned with GFP. White scale bar indicates 10, 5, 5, 2 μm in conidia, germination tube, appressorium and hyphae tip.

**Figure S3. MoSpa2 organizes actin nucleation and keep nucleation centre at hyphae tip. (related to Fig. 2).**

**A.** Actin filaments signal indicated by ABP140-3GFP in wild type (WT) and *SPA2* knockout mutant (*spa2-null*) of *Saccharomyces cerevisiae*. Black scale bar indicates 5 μm.

**B.** Quantification data of actin signal mean intensity per filament in (A). mean ± SE. Student *t*_test was used for statistical analysis, no significant difference showing between WT and mutant.

**C.** Actin profile of *M. oryzae* vegetative hyphae tip within 1.326 μm. Representative images of each strain are listed at right side. White bar cross tips means area for gray intensity profile production. White scale bar indicates 1 μm.

**D.** Living-cell images with time lapse of *M. oryzae* vegetative hyphae growth speed. White scale bar indicates 10 μm.

**E.** Quantitative results based on (D). One-way anova was used for statistical analysis relative to WT. ** means *p* < 0.01. Value = mean ± se. n = 9, 13, 5, in wild type, Δ*spa2-7*, Δ*spa2-6*.

**F.** Super resolution images with z-stack (0.2 μm as interval) of vegetative hyphae tip. 100 nM LatB was applied for treatment 20 minutes. AW means after washing of LatB 20 minutes. Green channel means Lifeact-GFP, red channel means FM4-64 dye, merge channel show overlap of SPK centre and actin nucleation centre. White arrow indicates SPK centre stained by FM4-64, red arrow means actin nucleation centre. n = 22, 11, 15, 7 in wild type, Δ*spa2-7*, Δ*spa2-6* and complementary line. White scale bar indicates 10 μm.

**Figure S4. Protein structure, purity and sequence characteristics. (related to Fig. 3).**

**A.** Predicted Alphafold based protein structure of MoSpa2, purple blue: 1-338 aa, red: 339-747 aa, cycan: 748-951 aa.

**B.** Protein sequence characteristics of MoSpa2 by MolPhase.

**C.** Purified recombinant proteins on SDS-PAGE gel. Stars indicate the target size.

**D-G.** Phase images of MoSpa2(1-747), MoSpa2(1-338), MoSpa2(339-747) and MoSpa2(748-951) for phase diagram generation. White scale bar in broad view images indicate 5 μm, and 1 μm in zoom images.

**A. H.** Phase diagram of MoSpa2(1-747), MoSpa2(1-338), MoSpa2(339-747) and MoSpa2(748-951) cross protein dose and ionic strength.

**I.** Representative FRAP images of 5 μM MoSpa2(1-747), MoSpa2(1-338) and MoSpa2(339-747), MoSpa2(748-951), containing 10% Alex-647 labelled proteins under 150 mM NaCl. White circle indicates fluorescence bleach region. White scale bar means 1 μm.

**J-K** Quantification FRAP results of (I). n = 12, 9, 12, 12, 17 in MoSpa2(1-747), MoSpa2(1-338), MoSpa2(339-747) and MoSpa2(748-951). One way anova with Sidak’s multiple comparisons test was used for statistical analysis between MoSpa2(1-747) and MoSpa2(1-338), MoSpa2(339-747), MoSpa2(748-951). * means *p* < 0.05. ns means no significant difference.

**Figure S5. MoSpa2 has no obvious effects on actin polymerization progress. (related to Fig. 3).**

**A.** TIRF images of 20 nM MoSpa2 truncates alone with ATP-Actin. 0.5 μM ATP-Actin was used in this assay. Black bar indicates 10 μm.

**B.** Seeds number at 3 minutes with ROI at 1652 μm^2^ after polymerization initiation in (A). from left to right, n = 24, 7, 7, 7, 7, 7 ROIs from 2 independent tests. One way anova with multiple comparation test was used for statistical analysis relative to actin alone. ns means no significant difference.

**C.** Elongation rate of actin filaments in (A). Units were calculated based on 374 units per μm. from left to right, n = 28, 14, 14, 14, 14, 14 filaments cross 2 minutes. One way anova with multiple comparasion test was used for statistical analysis relative to actin alone. ns means no significant difference.

**D-H.** Pyrene assay curve of actin polymerization with MoSpa2 truncates without MoBni1FH1C. 2 μM ATP-Actin was used in this assay, every each curve denotes at least three biological replicates average.

**Figure S6. MoSpa2 coordinates with MoBni1FH1C for actin polymerization. (related to Fig. 3).**

**A.** Predicted Alphafold based protein structure of MoBni1, MoBni1FH1C indicated with 1002 to the end (1740 aa), coloured by light green and dark green. Green colour indicates MoBni1-Nter.

**B.** Protein sequence characteristics of MoBni1 based on MolPhase.

**C.** Protein purity and SDS page isolation result of MoBni1FH1C. Star indicates target protein size.

**D.** TIRF images with time lapse on actin polymerization with MoBni1FH1C and MoSpa2 truncates. Black bar indicates 5 μm.

**E.** Seeds number at 5 minutes with ROI at 361 μm^2^ after polymerization initiation in Figure 4F. from left to right, n = 24, 24, 24, 24, 24, 12, 12 ROIs. One way anova with multiple comparasion test was used for statistical analysis relative to actin alone. ** means *p* < 0.01.

**Figure S7. MoSpa2 at low concentration with MoBni1FH1C shows no obvious effects on actin polymerization. (related to Fig. 3).**

**A.** Pyrene assay curve of actin polymerization with MoBni1FH1C and MoSpa2 truncates. 2 μM ATP-Actin was used in this assay, every each curve denotes at least three biological replicates average.

**B.** TIRF images of MoBni1FH1C combined with 20 nM MoSpa2 truncates at ATP-Actin polymerization. 0.5 μM ATP-Actin was used in this assay. Black bar indicates 10 μm.

**C.** Quantification results on (B). Actin seeds numbers were recorded as per ROI at 361 μm^2^. from left to right, n = 24, 24, 12, 12, 12, 12, 12 ROIs from left to right. One way anova with multiple comparation test was used for statistical analysis relative to actin alone. ** means *p* < 0.01.

**D-E.** Representative single particle images of 10 nM MoBni1FH1C in the presence of 20 nM MoSpa2 truncates (D) or 2 μM MoSpa2 truncates (E). MoBni1FH1C labelled with Alex488 and MoSpa2 variants labelled with Alex647 under 150 mM NaCl. White bar indicates 5 μm.

**Figure S8. MoSpa2-N IDRs is required for actin filaments stabilization. (related to Fig. 4).**

**A.** TIRF images of 2 μM MoSpa2 truncates alone on ATP-actin polymerization. Black bar indicates 10 μm.

**B.** Actin filaments skewness based on 10 minutes time-lapse TIRF images with ROI at 457.28 μm^2^. Value denotes mean ± SD, n = 7, 8, 8, 9, 8, 8 ROIs in ATP-Actin, +MoSpa2(1-747), +MoSpa2(1-338), +MoSpa2(339-747), +MoSpa2(748-951), +MoSpa2(1-951).

**C, E.** TIRF images with time lapse of MoSpa2 truncates effects on ATP-actin depolymerization by MoCof1 (C) and ScCof1 (E). Black bar indicates 5 μm.

**D, F.** Quantification results of MoSpa2 effects on depolymerization. Data indicates average with SE from ROIs with 1048 μm^2^ (except 961 μm^2^ for AC+MoSpa2(1-951)) at 8, 9, 10, 9, 6, 10 in (D) from top to bottom. Data indicates average with SE from ROIs with 655.93 μm^2^ (except 570 μm^2^ for AC+ScSpa2(1-951)) at 16, 10, 10, 9, 10, 7 in (F) from top to bottom.

**A. G.** Time lapse TIRF images of 4 μM MoSpa2(748-951) effects on anti-inhibition of MoSpa2(1-338) and MoSpa2(1-747)-induced inhibition on 100 nM MoCof1-drived actin filaments depolymerisation. Black scale bar indicates 5 μm.

**B. H.** Quantification results of MoSpa2(748-951) anti-inhibition of MoSpa2(1-338) and MoSpa2(1-747) on depolymerization. Data indicates average with SE from ROI with 447 μm^2^ (1047 μm^2^ for MoSpa2(748-951)+MoCof1) at 10, 8, 10, 10, 12, 12 from top to down samples.

**Figure S9. WH2 sequence alignment suggests MoSpa2(1-338) contains WH2-like sequence (related to Fig. 4).**

**A.** Sequence alignment of well-known WH2 sequences with MoSpa2(1-338) and ScSpa2(1-281). Sequences were submitted into Cluster-w for blast and Jalview is used for sequence alignment analysis.

**B.** WH2-like peptide within MoSpa2(1-338) and its structure with Actin binding model by Alphafold. Sequence highlighted with Cyan denotes helix while red means disordered region. Actin is indicated with green colour.

**C.** Colocalization of ScSpa2(1-281) and actin filaments in vitro. ScSpa2(1-281) at 4 μM was mixed with 10% ScSpa2(1-281)-Alex488 labelled protein, and inoculated with Actin filaments that polymerized from 2 μM G-actin and diluted 60 times. F-actin was stained with final concentration of 20 nM Phallodin565 for visualisation. White scale bar indicates 5 μm.

**Movie S1. MoSpa2 regulates actin assembly for hyphae vegetative growth.**

Source images with time lapse (4 minutes in total, 30 seconds interval) for Figure S3D. White scale bar indicates 5 μm.

**Movie S2. FRAP progress on MoSpa2 truncates undergo LLPS.**

Source images of Figure S4I. White scale bar indicates 1 μm. Fluorescence was bleached before 4 frame of images collection (1 second at interval), every 15 seconds was recorded for fluorescence recovery with 5 minutes duration. From left to right, MoSpa2(1-747), MoSpa2(1-338) and MoSpa2(339-747), MoSpa2(748-951)..

**Movie S3. MoSpa2 shows no significant induced actin polymerization activation.**

Source images with time lapse (7 minutes duration with 15 seconds interval) for Figure S5A. Scale bar indicates 5 μm. All MoSpa2 truncates at 20 nM and 0.5 μM G-actin.

**Movie S4. MoSpa2 induces MoBni1 activation for actin polymerization.**

Source image with time lapse (10 minutes duration with 15 seconds interval) for Figure S6D. Scale bar indicates 10 μm. All MoSpa2 truncates at 2 μM and MoBni1FH1C at 2.5 nM, 0.5 μM G-actin.

**Movie S5. Non-phase dose of MoSpa2 has no effects on increased MoBni1 activation.**

Source image with time lapse (10 minutes duration with 15 seconds interval) for Figure S7B. scale bar indicates 10 μm. All MoSpa2 truncates at 0.02 μM and MoBni1FH1C at 2.5 nM, 0.5 μM G-actin.

**Movie S6. MoSpa2 inhibits MoCof1-drived actin filaments depolymerization.**

Source images with time lapse (3.45 minutes duration with 15 seconds interval) for Figure S8C. Red scale bar indicates 5 μm. All MoSpa2 truncates at 4 μM, MoCof1 at 100 nM and G-actin at 0.5 μM.

**Movie S7. MoSpa2 inhibits ScCof1-drived actin filaments depolymerization.**

Source images with time lapse (4.45 minutes duration with 15 seconds interval) for Figure S8E. Red scale bar indicates 10 μm. All MoSpa2 truncates at 4 μM, ScCof1 at 100 nM and G-actin at 0.5 μM.

**Movie S8. MoSpa2(748-951) inhibits MoSpa2(1-338)-drived inhibition of MoCof1-drived actin filaments depolymerization.**

Source images with time lapse (5 minutes duration with 15 seconds interval) for Figure S8G. Scale bar indicates 5 μm. All MoSpa2 truncates at 4 μM, MoCof1 at 100 nM and G-actin at 0.5 μM.

